# Effects of background and contour luminance on the hue and brightness of the Watercolor effect

**DOI:** 10.1101/223792

**Authors:** Peggy Gerardin, Michel Dojat, Kenneth Knoblauch, Frederic Devinck

## Abstract

Conjoint measurement was used to investigate the joint influences of the luminance of the background and the inner contour on hue- and brightness filling-in for a stimulus configuration generating a water-color effect (WCE), i.e., a wiggly bi-chromatic contour enclosing a region with the lower luminance component on the exterior. Two stimuli with the background and inner contour luminances covarying independently were successively presented, and in separate experiments, the observer judged which member of the pair’s interior regions contained a stronger hue or was brighter. Braided-contour control stimuli that generated little or no perceptual filling-in were also used to assess whether observers were judging the interior regions and not the contours themselves. Three nested models of the contributions of the background and inner contour to the judgments were fit to the data by maximum likelihood and evaluated by likelihood ratio tests. Both stimulus components contributed to both the hue and brightness of the interior region with increasing luminance of the inner contour generating an assimilative filling-in for the hue judgments but a contrast effect for the brightness judgments. Control analyses showed negligible effects for the order of the luminance of the background or inner contour on the judgments. An additive contribution of both components was rejected in favor of a saturated model in which the responses depended on the levels of both stimulus components. For the hue judgments, increased background luminance led to greater hue filling-in at higher luminances of the interior contour. For the brightness judgments, the higher background luminance generated less brightness filling-in at higher luminances of the interior contour. The results indicate different effects of the inner contour and background on the induction of the brightness and coloration percepts of the WCE, suggesting that they are mediated by different mechanisms.

## 1. Introduction

Color appearance is not determined only by the local light signals from each object but is also influenced by global contextual features. The watercolor effect (WCE) is an interesting phenomenon for studying such processes (Pinna, 1987; Pinna et al., 2001). A pair of wiggly contours composed of a light chromatic contour (e.g., orange) surrounded by a darker chromatic contour (e.g., purple) bounding an achromatic surface area elicits a filling-in of the hue of the lighter contour over the entire enclosed area (Figure 1a). The WCE is distinguished from other assimilation illusions by its large spatial extent; the phenomenon has been observed over distances of up to 45 deg (Pinna et al., 2001). In addition to the assimilative color spreading, the subjectively colored area is perceived as figure while the surrounding area appears as ground (Pinna et al., 2003; Pinna & Tanca, 2008; Tanca & Pinna, 2008).

**Fig. 1:**
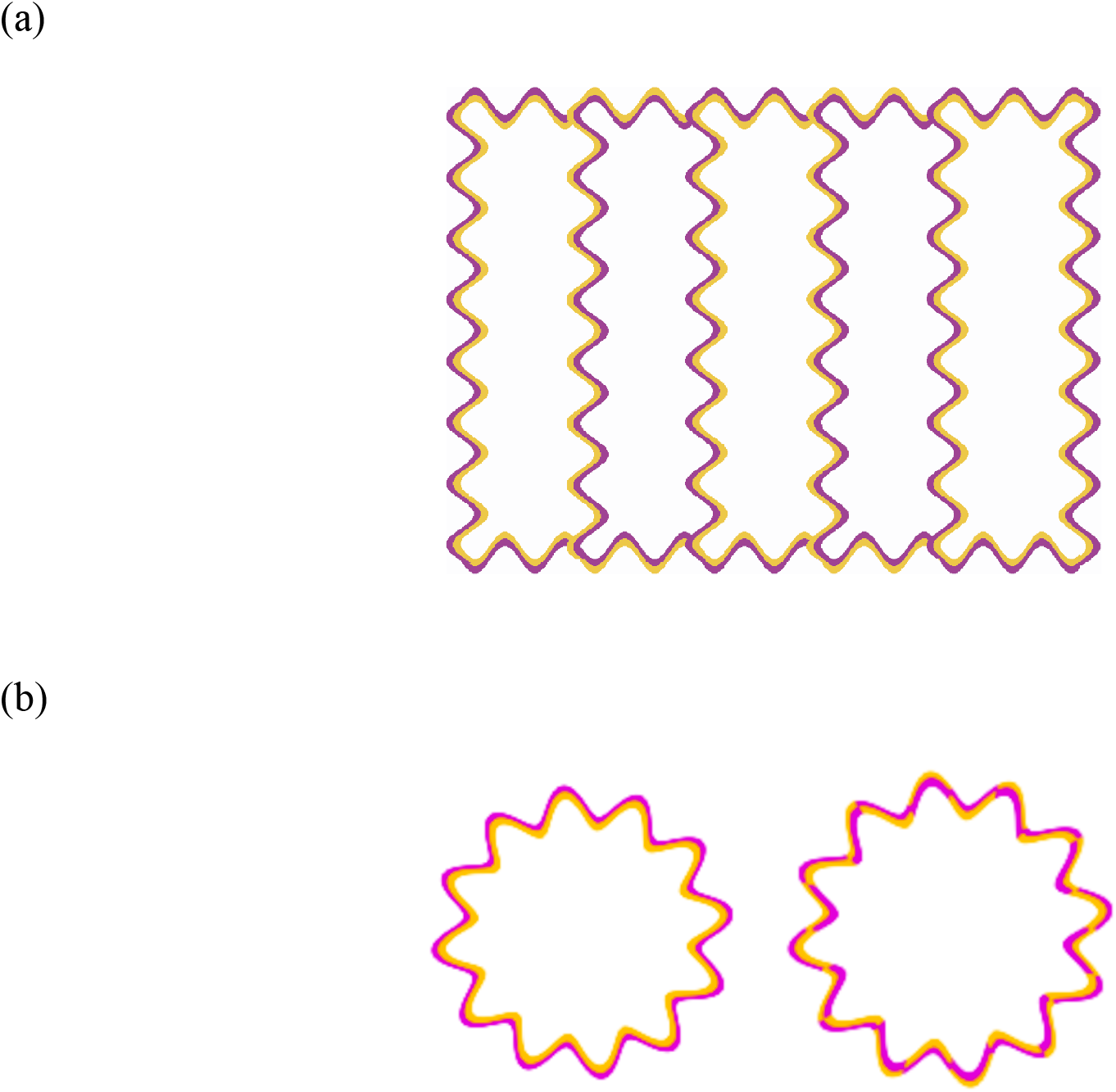
(a) Example of the Watercolor Effect. When a light orange contour is surrounded by a dark purple contour, the enclosed area takes the tint of the orange border. (b) Example of stimuli using Fourier descriptor as test stimulus (presented on the left side) and using braided contour as control stimulus (displayed on the right side).

Studies of the WCE have typically examined the effects of the inducer configuration producing the WCE. For example, the intensity of the filling-in percept appears greater with increases in luminance contrast between the inner and outer contours for an achromatic WCE (Cao et al., 2011) and for a WCE that has both luminance and chromatic components (Devinck et al., 2005; Devinck & Knoblauch, 2012). Devinck et al. (2005) noted that observers did not need to modify significantly the luminance of the enclosed area in a matching experiment. Other critical characteristics of the inducing contours that modulate the strength of the WCE include the continuity and contiguity of the contour pairs (Devinck & Spillmann, 2009; Devinck & Knoblauch, 2012). Recent demonstrations of the sensitivity of the phenomenon to contour adaptation provide additional support for a role of contour integration mechanisms in the WCE (Coia & Crognale, 2017). The strength of the phenomenon was found to be size-tuned with the strongest WCE observed for a contour width of about 15 arcmin and was optimal for equal contour widths (Devinck et al., 2014a). While the WCE has been reported for linear contours (Pinna et al., 2001), its strength is nearly independent of the amplitude of contour undulation but increases with contour frequency up to an asymptotic level (Gerardin et al., 2014). Finally, Pinna et al. (2001) demonstrated that several different color pairs can generate the coloration effect (see also Devinck et al., 2005). Specifically, Devinck et al. (2006) demonstrated that the coloration effect is stronger when the chromatic contrast is larger. Thus, the coloration effect depends on a conjunction of chromatic and luminance contrasts but also on spatial parameters of the inner and outer contours.

The WCE is perceptually salient but has proved difficult to quantify with precision showing large variability within and across observers (Cao et al., 2011; Devinck et al., 2005; von der Heydt & Pierson, 2006). More recently, the WCE was quantified by using paired-comparison methods that have been extended to estimate perceptual scales within a signal detection framework (Devinck & Knoblauch, 2012). Two such procedures are Maximum Likelihood Difference Scaling or MLDS (Maloney & Yang, 2003; Knoblauch & Maloney, 2008, 2012) and Maximum Likelihood Conjoint Measurement or MLCM (Ho et al., 2008; Knoblauch & Maloney, 2012). Difference scaling is useful for measuring perceptual strength along a single physical dimension, whereas conjoint measurement was conceived to assess the combined effects of several dimensions on appearance (Falmagne, 1985; Knoblauch & Maloney, 2012; Krantz et al., 1971; Luce & Tukey, 1964; Roberts, 1979). MLCM has been successfully applied to estimate perceptual scales associated with different sets of physical continua including surface material properties (Ho, Landy & Maloney, 2008; Qi, Chantler, Siebert & Dong, 2015; Hansmann-Roth & Mamassian, 2017), color appearance (Gerardin et al., 2014; Rogers, Knoblauch & Franklin, 2016) and time perception (Lisi & Gorea, 2016). The signal detection decision model allows specifying the perceptual scales in terms of the signal detection parameter *d’* (Gerardin et al., 2014; Knoblauch & Maloney, 2012).

The aim of the present study is to estimate perceptual scales for two dimensions, the luminance elevation of the inner contour and the luminance elevation of the background. While the luminance contrast between the inner and outer contours has been tested intensively in the WCE, experiments evaluating the influence of the background luminance are scarce. Indeed, the WCE has generally been demonstrated for a background of higher luminance than both inner and outer contours. Although the surround (e.g., the background) is known to be an important influence of color appearance (Brenner & Cornelissen, 2002; Brown & MacLeod, ļ 997; Shevell, ļ 978; Walraven, ļ976), it has not been systematically explored for the coloration effect in the WCE. In addition, most studies of the WCE focus solely on its coloration effect. Here, we also investigate the influences of the background and inner contour luminances on the perceived brightness of the interior region. In summary, we employed conjoint measurement to study how both the background and the inner contour luminances influence judgments of both the hue and brightness in the WCE.

## 2. General Methods

### 2.1 Observers

Four observers participated in these experiments. Three were naïve and the fourth was one of the authors. Observers ranged in age between 26 and 40 years. All had normal color vision as tested with the Farnsworth Panel Dļ5, and had normal or corrected-to-normal visual acuity. Experiments were performed in accordance with the principles of the Declaration of Helsinki for the protection of human subjects.

### 2.2 Apparatus

Stimuli were presented on a NEC MultiSync FP2141sb color CRT monitor driven by a Cambridge Research ViSaGe graphic board with a color resolution of 14 bits per gun (Cambridge Research Systems, Rochester, United Kingdom). The experimental software was written to generate all stimuli, control stimulus presentation and collect responses in MATLAB 7.9 (MathWorks, http://mathworks.com), using the CRS Toolbox extensions. The monitor was calibrated using an OptiCal photometer with the calibration routines of Cambridge Resarch Systems. Observer position was stabilized by a chinrest and observer-to-screen distance was 80 cm. Experiments were performed in a dark room. Both eyes were used for viewing.

### 2.3 Stimuli

The stimuli were constructed as Fourier descriptors (Zahn & Roskies, 1972). Each stimulus was defined with respect to a circle of 3.2 deg diameter whose radius, *r*, was modulated sinusoidally as a function of angle according to the equation:

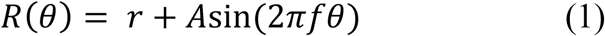

where *R* is the stimulus radius at angle *θ, r* the average radius of the stimulus, *A* the modulation and *f* the frequency in cycles per revolution (cpr). In the present study, the frequency was fixed at *f* = 10 cpr and the amplitude of both contours at *A* = 0.36 (Figure 1b, left).

All stimuli were composed of three colors: an orange inner contour (x,y = 0.44, 0.43) with the luminance varying from 30.02 cd/m^2^ to 62.74 cd/m^2^ and a purple outer contour (x,y = 0.3ļ, 0.ļļ; Y = 2ļ.ļ2 cd/m^2^), presented on a neutral white background (x,y = 0.29, 0.32) with the luminance of the background (both outside and inside of the contours) varying between 35.56 cd/m and 65.56 cd/m^2^. The contour pairs were each of width ļ6 arcmin, i.e., 8 arcmin for the inner and outer contours, each.

Stimuli were specified in the DKL color space (MacLeod & Boynton, ļ979; Krauskopf, Williams & Heeley, ļ982; Derrington, Krauskopf & Lennie, ļ984). DKL color space is a three-dimensional opponent-modulation space based on the Smith and Pokorny (ļ975) cone fundamentals. The sum of L and M cone excitations varies on one axis (luminance), while M cone excitation subtracted from L cone excitation varies on the second axis (L - M); and the sum of L and M cone excitations subtracted from S cone excitation varies on the third axis (S − (L + M)). The DKL axes were scaled between −ļ and ļ, where +/- ļ corresponds to the maximum contrast for each axis on the monitor. The stimuli were specified with the purple and orange contours at azimuth of 320 and 45 deg respectively. Luminance of the independent variables is specified as elevation from the equiluminant plane. The luminance elevations of the orange contour in DKL color space varied from −0.6 to 0 while the luminance elevation of the background ranged from −0.5 to 0.

In the present study, five levels of luminance inner contour and five levels of luminance background were used. All levels were crossed creating a 5 × 5 grid with a total of 25 stimuli. Figure 2 shows an example of the range of stimuli used, with the inner contour luminance varying across rows and the background luminance across columns.

**Fig. 2:**
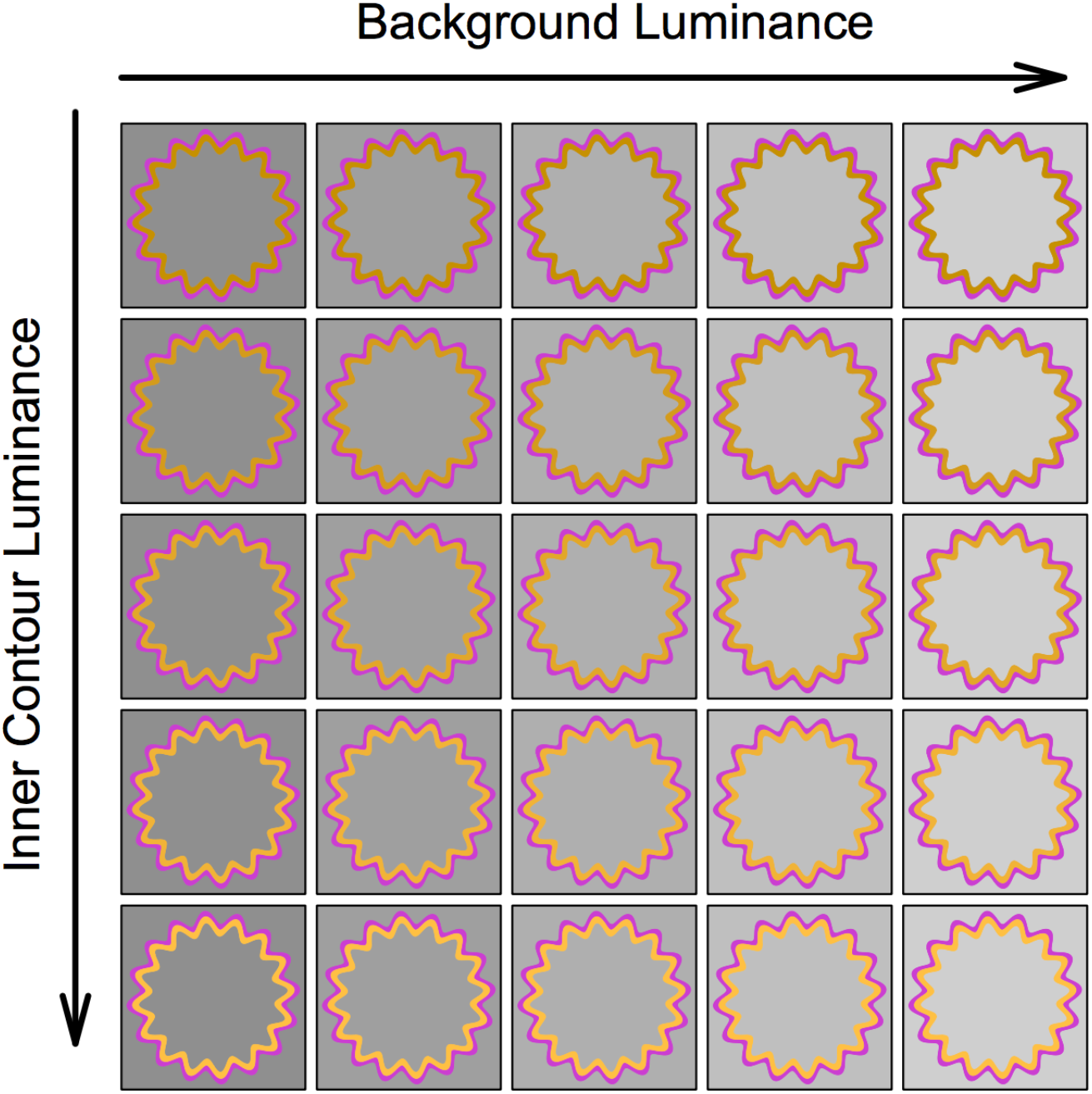
Examples of stimulus set used for a conjoint measurement experiment. The figure indicates the set of stimuli used in both judgment tasks. Each column corresponds to a different luminance elevation of the background and each row to a different luminance elevation of the inner orange contour.

Control stimuli were also tested consisting of patterns that were identical to the test stimuli except that the contours were interlaced (Figure 1b, right) and generated little filling-in, as previously demonstrated (Devinck & Knoblauch, 20ļ2; Devinck et al., 20ļ4a; Gerardin et al., 20ļ4). These control stimuli were used to verify that observers responded to the filling-in appearance and not to other stimulus features.

### 2.4 Procedure

On each trial, two different stimuli chosen randomly from the 5 × 5 grid were presented in succession to the observer. Observers performed two tasks in separate randomly ordered and counter-balanced sessions in which they compared the interior regions of the two successively presented stimuli. In the first task, observers were instructed to judge which central region evoked the strongest orange hue. In the second task, observers were asked to judge which central region appeared brighter. An equal number of test and control stimuli were interleaved in each session. With 5 levels along each of the dimensions varied, there are (25 * 24)/2 = 300 unordered pairs. Stimuli were randomly ordered for each presentation. On each trial, a randomly chosen pair of test or control stimuli was presented. A session consisted of the random presentation of all 600 test and control pairs. Each task was repeated five times, yielding 1500 test and 1500 control trials for each observer.

Prior to the experiment, observers were dark-adapted for 3 min. At the beginning of each trial, a fixation cross was presented in the center of the screen of duration 500 ms. At its extinction, the first pattern was presented during 500 ms followed by a fixation cross for 500 ms, and then the second pattern for 500 ms., followed by a blank screen. The observer’s response initiated a 1 s pause before the start of the next trial. An initial practice block of 10 trials preceded the experiment. The experimental session started, when the observer felt at ease with the task, otherwise additional practice sessions were run. A free viewing procedure was used to ensure that observer’s judgments were based on foveal views of the stimuli.

### 2.5 Model

The data were analyzed as a decision process within the framework of a signal detection model and fit by maximum likelihood (Ho et al., 2008; Knoblauch & Maloney, 2012). Three nested models of the decision process are fit to obtain the best prediction of the set of observers’ choices: an independence model, an additive model and a saturated model. Each model yields estimates of perceptual scale values or internal responses that have the property that equal differences in response are perceptually equal. The independence model fits the observer’s judgments based on only one of the component dimensions. The additive model fits the judgments based on the sum of component psychological responses generated by the physical dimensions. The saturated model fits the observer’s judgments including an interaction term that depends on the specific levels of the two components in addition to their simple additive combination. The three models are then evaluated using a nested likelihood ratio test. This is done separately for the experiments based on hue, and brightness judgments. The formal description of the model is described next and follows similar descriptions elsewhere (Gerardin et al., 20ļ4; Ho et al., 2008; Knoblauch & Maloney, 20ļ2).

We represent the stimulus levels along the two dimensions by a variable *φ_i,j_*, where *i* and *j* correspond to the luminance levels of the background and the inner contour, respectively. In the decision models, each of the dimensions contributes a response, 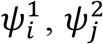, to the intensity of the perceived filling-in depending on the corresponding physical intensity levels, where the superscripts correspond to the responses to the background and interior contour luminances, respectively. In the additive model, when observers judge which central area is the more saturated orange color or which appears to be brighter, we suppose that I the filling-in response is the sum of the component responses

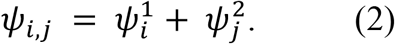

Observers compare the two presented central surfaces and the difference between the filling-in strength of the stimulus *φ_i,j_* and the stimulus *φ_k,l_* is computed as follows

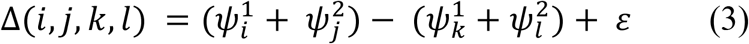

where *ε* refers to additive noise in the decision process and is modeled as a Gaussian random variable with *μ =* 0 and variance = *σ*^2^. In plots, we indicate the stimulus level by the index and not by the physical units, allowing both dimensions to be plotted together. With 5 levels along each dimension, there are 2 * 5 levels plus 1 variance = 11 parameters to estimate. To make the model identifiable, however, the response at the lowest level along each dimension is arbitrarily set to 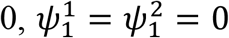, and the variance is fixed to 1 for each estimated value, yielding only 8 parameters to estimate. The parameter values, 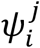 are chosen to maximize the likelihood, 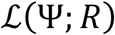, over the ensemble of choices, *R*, made by the observer.

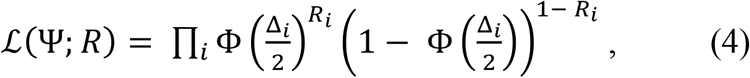

where Φ is a cumulative distribution function for a Gaussian with mean 0 and variance 1. The 2 in the denominator of the argument scales the variance for each value of *ψ* to 1 so that the perceptual scale values are parameterized in terms of *d’*. In practice, this is performed using a Generalized Linear Model with the MLCM package in R (Knoblauch & Maloney, 20ļ2, 20ļ4).

If the observer’s judgments depend on only one of the component dimensions, we obtain the independence model, reducing the decision variable to

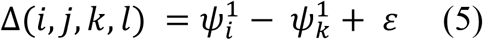

where the judgments depend on only dimension ļ, here. In this reduced model, the values of 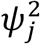 are fixed at 0 and there are only 4 free parameters to estimate. Replacing the superscript ļ by 2 yields the independence model for the other dimension.

Finally, the saturated observer model includes an interaction factor that depend on the intensity levels of both dimensions; the decision variable is defined as follows:

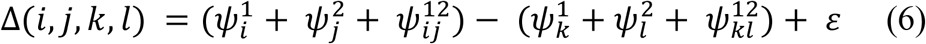

Due to the interaction terms, the responses cannot be explained by a simple additive combination of components as in the previous two models. With 5 levels along each dimension, only one cell in the 5x5 grid of responses is fixed at 0 leading to 24 (the maximum) free parameters to estimate, which is the origin of the term saturated.

We analysed the data using the MLCM package (Knoblauch & Maloney, 20ļ2, 20ļ4) in the open source software R (R Core Team, 20ļ7) to estimate the perceptual scale values and model the contribution of both dimensions. The likelihood ratio tests were evaluated using a *χ*^2^ statistic with degrees of freedom the difference in number of parameters fit for each pair of models.

## 3. Results

Judgments based on color and on brightness from the observers are shown in Figure 3 for test and control conditions in Conjoint Proportion Plots or CPP (Ho et al., 2008; Knoblauch & Maloney, 20ļ2). In the CPP, the raw data are presented in a grid format in which each cell of the grid corresponds to one stimulus pair comparison. Each CPP contains all stimulus combinations and summarizes the proportion of times the stimulus *S_kl_* was judged for one response criterion, hue (a) or brightness (b), to show a greater filling-in than the stimulus *S^ij^*, coded according to the grey levels indicated by the color bar presented on the right side of each graph. The levels of both dimensions are represented along each axis where the 5×5 outer check indicates the stimulus levels along one dimension and with each outer check subdivided into smaller 5×5 checks indicating the stimulus levels for the second dimension. Figure 3c shows the expected pattern of responses for an ideal observer who chooses only the higher level along one of the two stimulus dimension. The CPP presented on the left side indicates the results when the judgments depend on the first dimension alone (here, the background luminance) and the CPP displayed on the right side when the judgment depend on the second dimension (here, the inner contour luminance).

Results from the hue and brightness judgments for each observer are shown in Figure 3a and 3b respectively with the results displayed on the top row for the test condition and on the bottom row for the control condition in each figure. For the hue judgments, the CPP for the test stimuli resembles more closely the ideal CPP displayed for the second dimension, suggesting that the luminance of the inner contour contributed more strongly to the choices than the luminance of the background. For the brightness judgments, the CPP for the test stimuli is more similar to the ideal CPP displayed for the first dimension, indicating that the luminance of the background contributed more strongly to the choices in comparison with the luminance of the inner contour. Deviations from the ideal patterns, however, indicate contributions from both dimensions for both tasks.

In these experiments, observers were instructed to judge the appearance of the interior region of the stimulus. However, it is possible that observers attended to the experimental dimensions (e.g., the continuity of the color of the contour) instead of the appearance of the interior region. If this were the case, we should obtain the same pattern of responses between the test and the control conditions. However, the test CPP patterns are different, with the control CPP patterns showing little, if any, systematic structure.

**Fig. 3:**
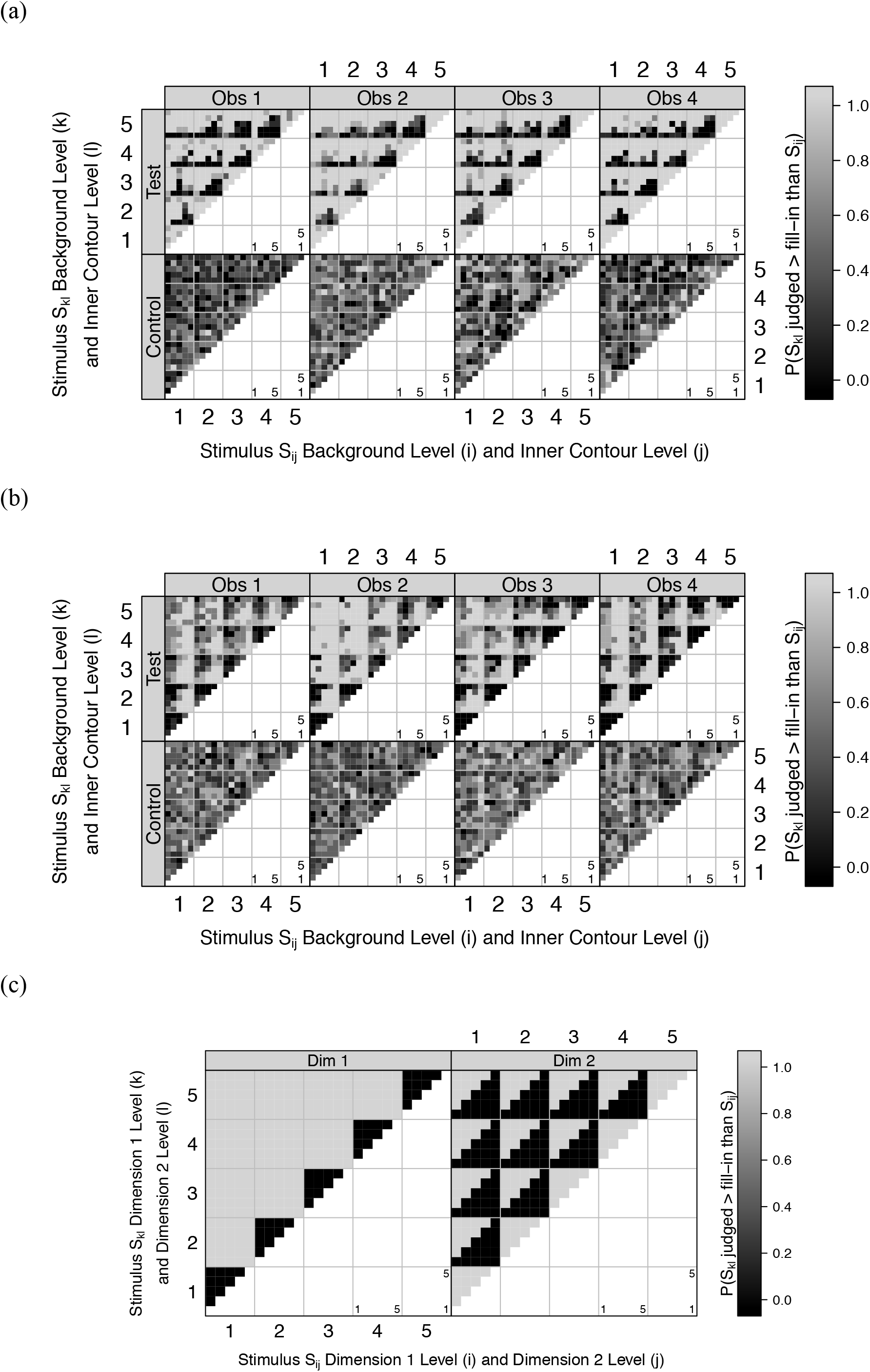
Conjoint proportion plots for judgments based on hue (a) and on brightness (b). Each plot shows the proportion of stimulus *S* judged to have a greater filling-in than the stimulus represented in abscissa *S^ij^* as grey level according to the color bar on the right side. The luminance elevation of the background is indicated by the large grids (*i,k*) and each grid is subdivided into smaller 5×5 grid indicating the luminance elevation of the inner contour (*j,l*). In each set of graphs, the top row indicates the results for the test stimuli and the bottom for the control stimuli for 4 observers. (c) Conjoint proportion plots for a simulated observer. Expected responses for an observer who judges the stimuli based on the contributions along only one of the dimensions.

Statistical comparisons of additive and independence models based on a nested likelihood ratio test are shown in Table I for judgments based on hue and in Table II for judgments based on brightness. In these tables, each row indicates the test for an observer, and the last column corresponds to the probability that the additive model fits no better than the independence model. The degrees of freedom are obtained from the difference of the number of coefficients estimates in each model (8 (additive) – 4 (independence) = 4). Comparing the independence with the additive model indicates that the independence model can be rejected for the test stimuli for both tasks and for all observers *(p* < 0.001). The motivation for testing the nested decision models for the control stimuli is less clear. Instead, we used a linear mixed-effects model to test if the estimated perceptual scale values depended significantly on the stimulus level with a random observer intercept (Bates et al., 2015). For both the hue and the brightness judgments, no significant dependence was found (hue: *χ*^2^(30) = 38.5, *p* = 0.14; brightness: *χ*^2^(30) = 22.2, *p* = 0.85). It could be argued that 4 observers is not sufficient to estimate the variances accurately in a mixed-effect model. We also performed the tests using a linear model with Observer entering as a fixed-effect, with no significance obtained in either case. The advantage of the mixed-effects model is that the results generalize to the population rather than just the sample of 4 observers tested (Knoblauch & Maloney, 20ļ2; Moscatelli et al., 20ļ3; Pinheiro & Bates, 2000).

**Table I:** Comparison of independence and additive models for judgements based on hue

|  | Test stimuli (additive model × independent model) |  |  |
| --- | --- | --- | --- |
|  | Df | Deviance | p-value |
| Obs. 1 | 4 | 270.22 | < 0.001 |
| Obs. 2 | 4 | 568.41 | < 0.001 |
| Obs. 3 | 4 | 309.60 | < 0.001 |
| Obs. 4 | 4 | 406.38 | < 0.001 |

**Table II:**
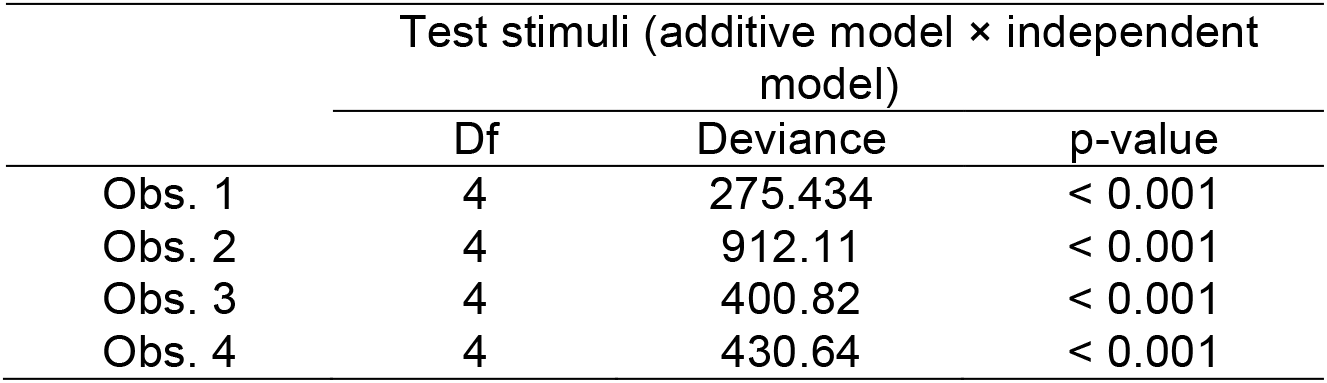
Comparison of independence and additive models for judgement based on brightness

|  | Test stimuli (additive model × independent model) |  |  |
| --- | --- | --- | --- |
|  | Df | Deviance | p-value |
| Obs. 1 | 4 | 275.434 | < 0.001 |
| Obs. 2 | 4 | 912.11 | < 0.001 |
| Obs. 3 | 4 | 400.82 | < 0.001 |
| Obs. 4 | 4 | 430.64 | < 0.001 |

Figure 4 shows the estimated contributions of each dimension obtained from fitting the additive model for each of the tasks. For judgments based on hue, the average estimated scales for each pair of inner contour luminance and background luminance elevation are shown in Figure 4 (a) for 4 observers. In this figure, the column labels indicate the observer identification. The top row shows the scale values estimated for the test stimulus and the bottom for the control stimulus. Black circles indicate the inner contour contribution and white the background contribution to the judgments. The values on the abscissa indicate the five stimulus levels for each dimension coded from ļ to 5. These values are indices to the 5 luminance elevations of the inner contour and the background used in the experiment.

The results in Figure 4a indicate that the contribution of the luminance of the interior contour dimension to the hue filling-in strength of the inner contour as does the contribution of the background increases with luminance elevation but with a smaller effect. The second row shows results obtained for the control stimuli. Here, there appears no systematic influence of either dimension on the strength of the coloration effect, because both dimension contributions are close to zero at all stimulus levels. This result further supports that the observers based their judgments on the perceived filled in color of the interior rather than the luminance of the inner contour or the background.

**Fig. 4:**
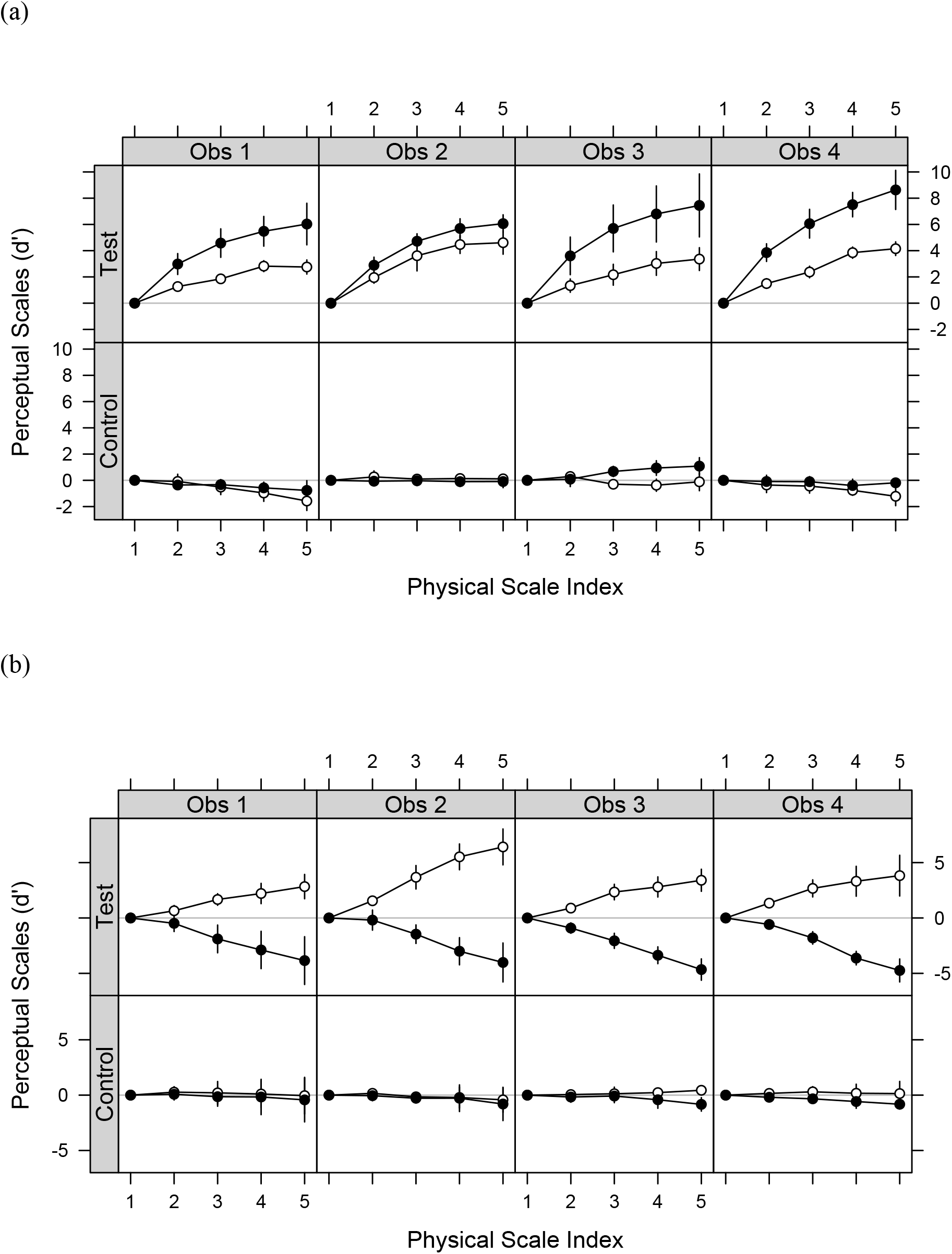
(a) Estimated scales for judgments based on hue. Additive model average estimates for test (top row) and control stimuli (bottom row) as a function of inner contour elevation (black circles) and luminance background elevation (white circles) for four observers. Error bars show 95% confidence intervals for estimates across the 5 runs. (b) Results for judgments based on brightness. The solid lines indicate the estimated contributions of each dimension under the additive model for test (top row) and control patterns (bottom row) as a function of inner contour elevation (black circles) and luminance background elevation (white circles) for four observers. Error bars show 95% confidence intervals for estimates across the 5 runs.

Figure 4 (b) shows the average estimated scales for each observer for each pairing of inner contour and background luminance elevations when judgments were based on the brightness of the interior region. The information in the figure is organized in the same fashion as Figure 4a. The top row shows the additive model fits to judgments of the brightness of the interior region. As the inner contour increases in luminance, the estimated contribution for this component decreases, indicating that the brightness of the interior region decreases, i.e., generating a relative contrast rather than an assimilative effect. The background dimension contributes positively with the luminance elevation to judgments of brightness. These results contrast with the scales estimated for the control stimuli that show no difference and little effect of both dimensions on the strength of brightness filling-in.

Temporal forced choice experiments can be subject to order effects (Yeshurun et al., 2008). To test for this, we compared the scales obtained for trials in which the higher luminance was presented in the first interval with the trials in which it was presented in the second interval. We performed the same analysis with respect to the luminance of the inner contour. For the hue judgments, the average estimated scales for each background luminance order are shown in Figure 5 (a) for 4 observers, and judgments based on brightness are displayed in Figure 5 (b). In these figures, the column labels indicate the observer identification. The top row shows the estimated scale values for the contribution of the inner contour luminance and the bottom for the background dimension. White circles indicate the dimension contribution to the judgments when the luminance elevation of the background is higher in the first stimulus presentation and black circles the contribution to the judgments when the luminance elevation of the background is higher in the second presentation. The values on the abscissa indicate the five levels of each dimension coded from ļ to 5. For judgments based on hue, the results in Figure 5a indicate that the contribution of the inner contour dimension is not different between both background presentation orders. Moreover, the background dimension is approximately the same between the presentation orders. Similar results were obtained when judgments were based on brightness (Figure 5b). This shows that any order effect based on the luminance background is very slight or absent in our experiments.

For completeness, we show in Figure 6 the effect of the presentation order of the luminance of the inner contour on the estimated scales for both judgments organized in the same fashion as Figure 5. For both tasks, the scales for neither dimension were influenced as a function of the presentation order.

**Fig. 5:**
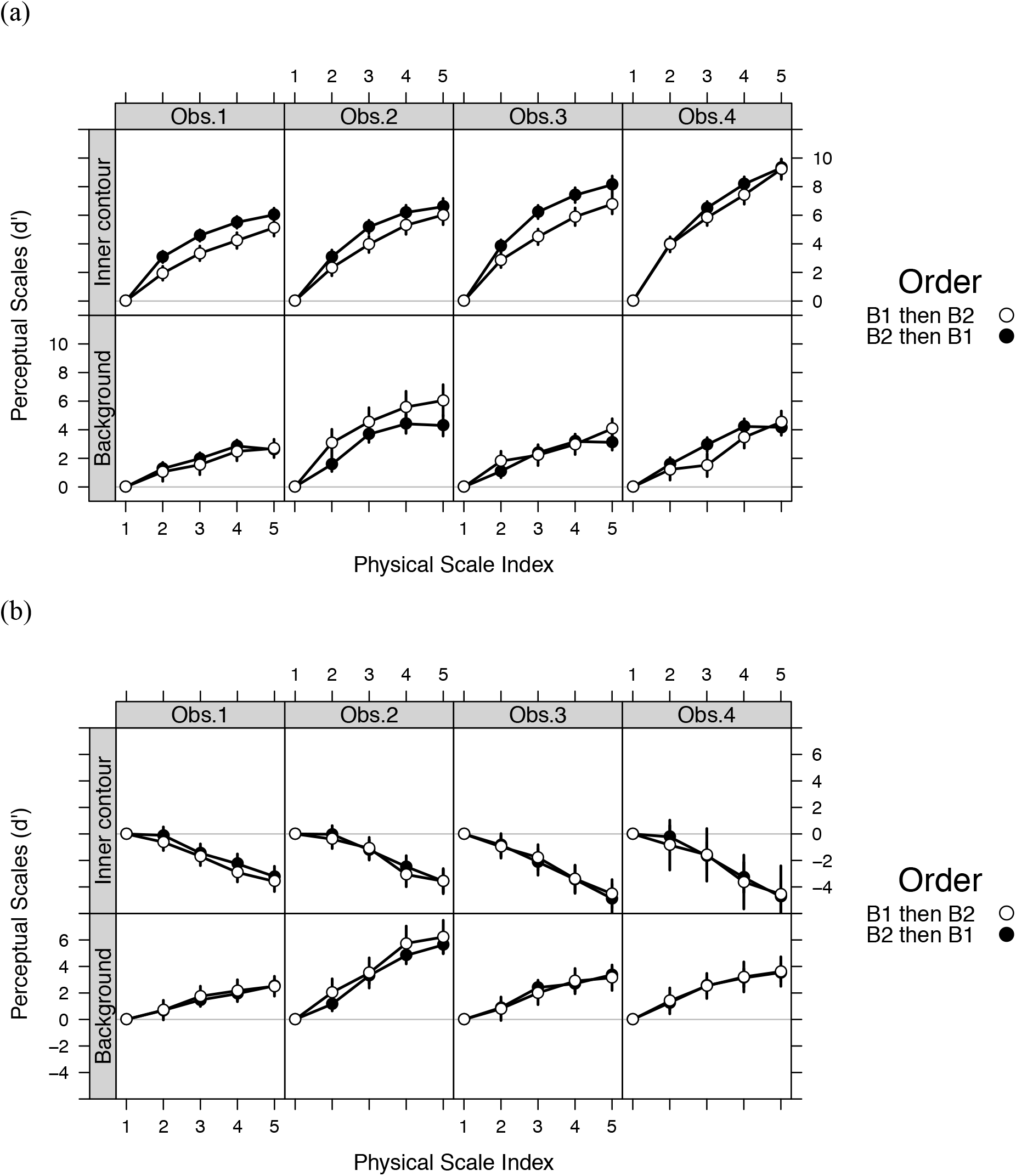
(a) Results for judgments based on hue depending on the presentation order for the luminance background of the test stimulus. The solid lines indicate the estimated contributions under the additive model for the inner contour dimension (top row) and the background dimension (bottom row) as a function of the background order presentation. The white circles are used to indicate that the luminance elevation of the background is higher in the first interval than in the second interval and the black circles are used to represent a higher luminance elevation of the background in the second interval than in the first interval. Error bars are 95% confidence intervals based on a bootstrap procedure (Knoblauch & Maloney, 2012). (b) Results for judgments based on brightness depending of the luminance background presentation order for the test stimulus. Additive model average estimates for the inner contour dimension (top row) and the background dimension (bottom row) when the luminance elevation of the background is higher in the first interval than in the second interval (white circles) and when the luminance elevation of the background is higher in the second interval than in the first interval (black circles). Error bars are 95% confidence intervals based on a bootstrap procedure (Knoblauch & Maloney, 2012).

**Fig. 6:**
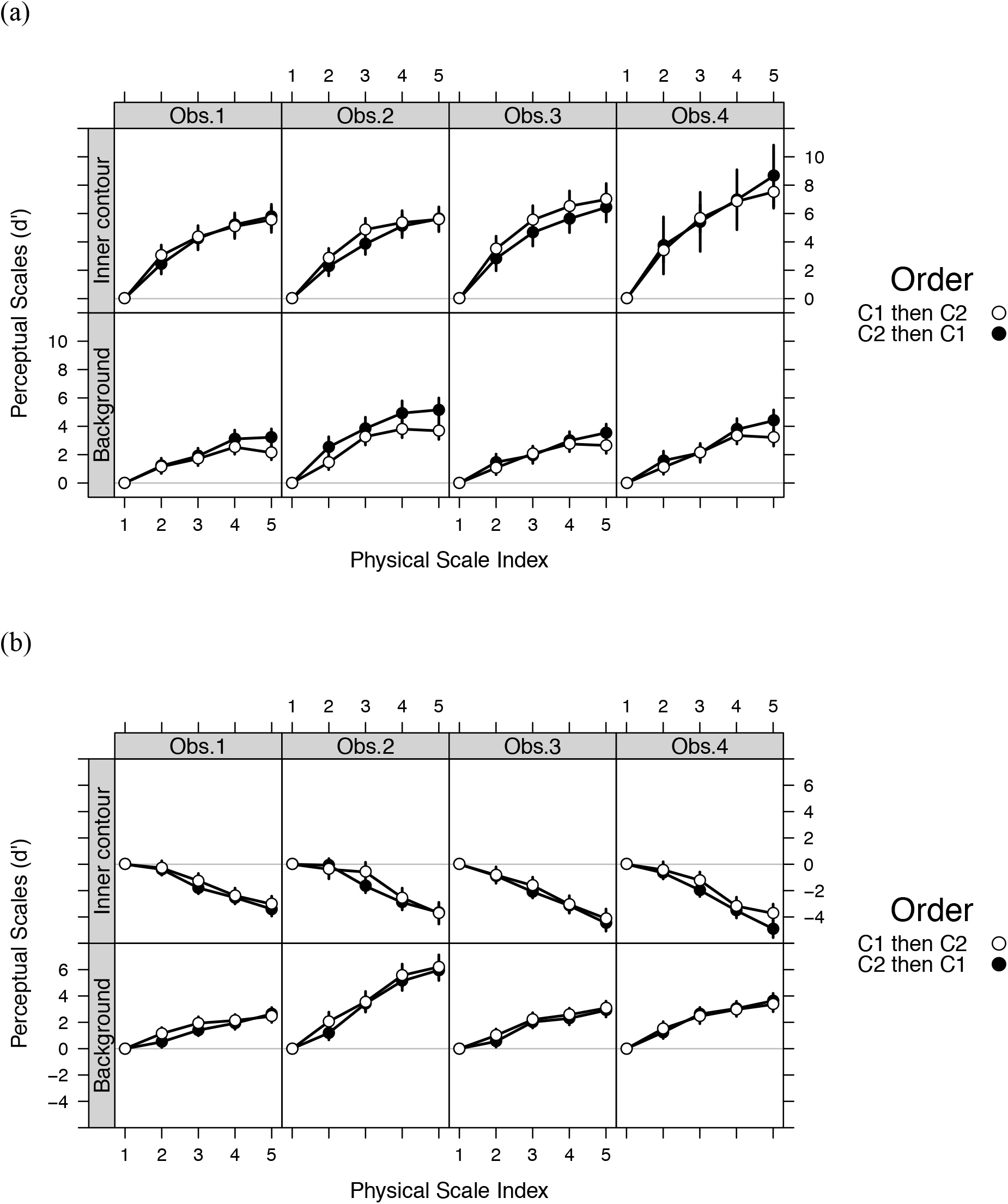
Results for judgment based on hue (a) and on brightness (b) depending on the presentation order for the luminance contour of the test stimulus. The information in the figure is organized in the same fashion as Figure 5. The white circles are used to indicate that the luminance elevation of the inner contour is higher in the first interval than in the second interval and the black circles are used to represent a higher luminance elevation of the inner contour in the second interval than in the first interval. Error bars are 95% confidence intervals.

Comparing the saturated and additive models with nested likelihood ratio tests rejected the hypothesis that the saturated model provided a fit no better then the additive model, thus, demonstrating that an interaction term is required to describe the observers’ judgments for both tasks (Tables III and IV). The degrees of freedom indicate the difference of the number of coefficents estimated in the 2 models (24 (saturated) − 8 (additive) = ļ6). The additive model fit was rejected in all 8 tests. The estimated coefficients for the saturated model are shown for each observer and both tasks in the panels of Figure 7. In these displays, the estimated scale values are plotted as a function of the stimulus index for the luminance of the inner contour with the index of the background luminance specified as a parameter for each curve. The results for the hue judgments are presented on the top row with the brightness judgments on the bottom. The averages of the four observers for both judgment conditions are summarized as three-dimensional surfaces in Figures 8a and b. If the additive model was a good fit to the data, the curves for different background levels would be parallel in Figure 7. However, for both the hue and brightness responses, as the background luminance increases, the curves fan out. For the hue judgments, this results in a larger range of hue responses to the range of inner contour luminances tested for the higher than lower luminance backgrounds. For the brightness judgments, in contrast, the range of response decreases at the highest luminance background. Thus, the background luminance produces different effects on both the type of filling-in (assimilation vs contrast) for hue and brightness judgments and on the dynamic range of the response, expanding it for hue judgments but compressing it for brightness.

**Fig. 7:**
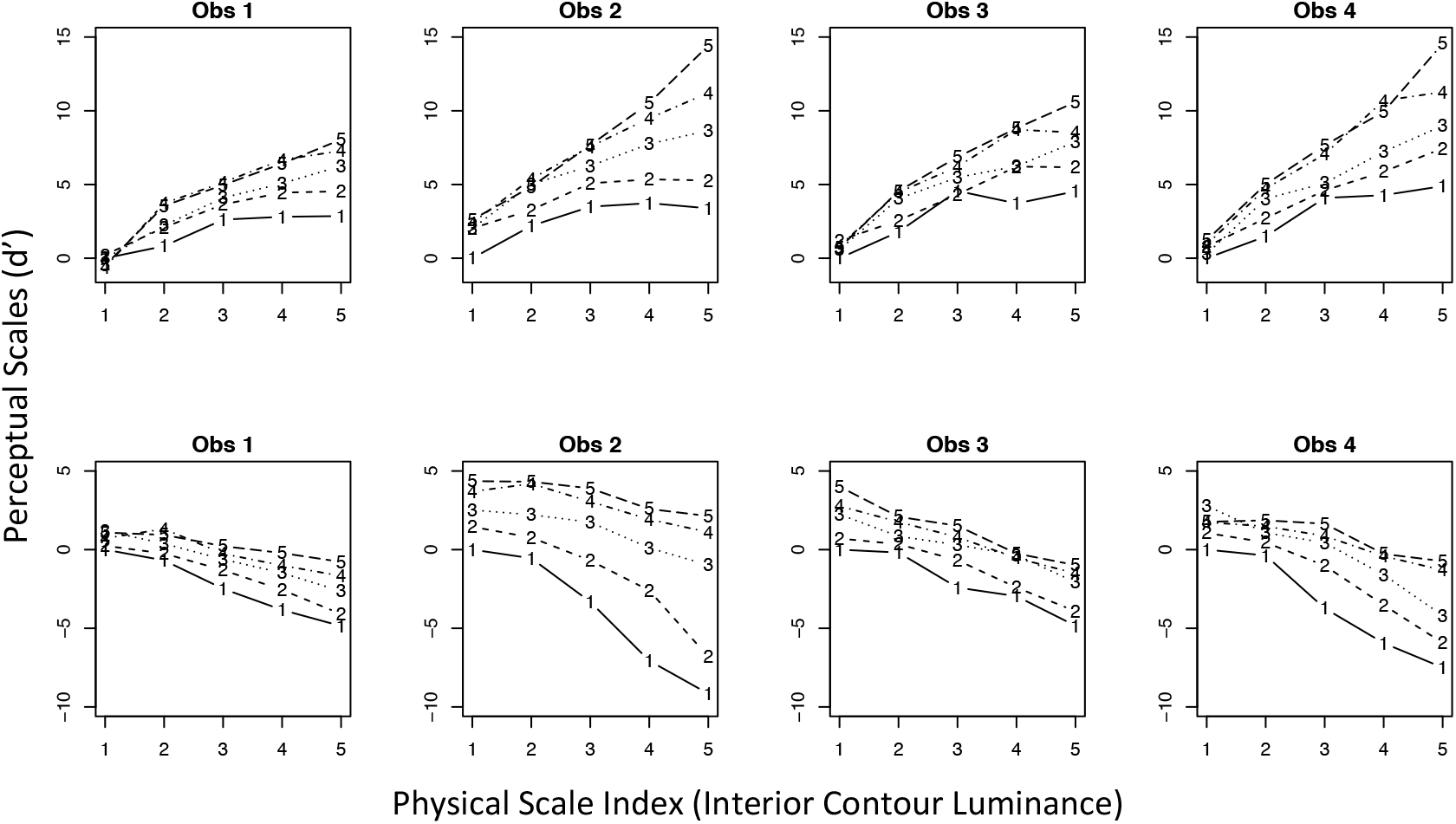
Results of the estimated contributions for each combination of the two dimensions under the saturated model for four observers. The different lines are used to code the index of the background dimension (for indices 1 to 5). The top row represents the estimated contribution when judgment is based on hue and the bottom row is the estimated contribution when judgment is based on brightness.

**Fig. 8.**
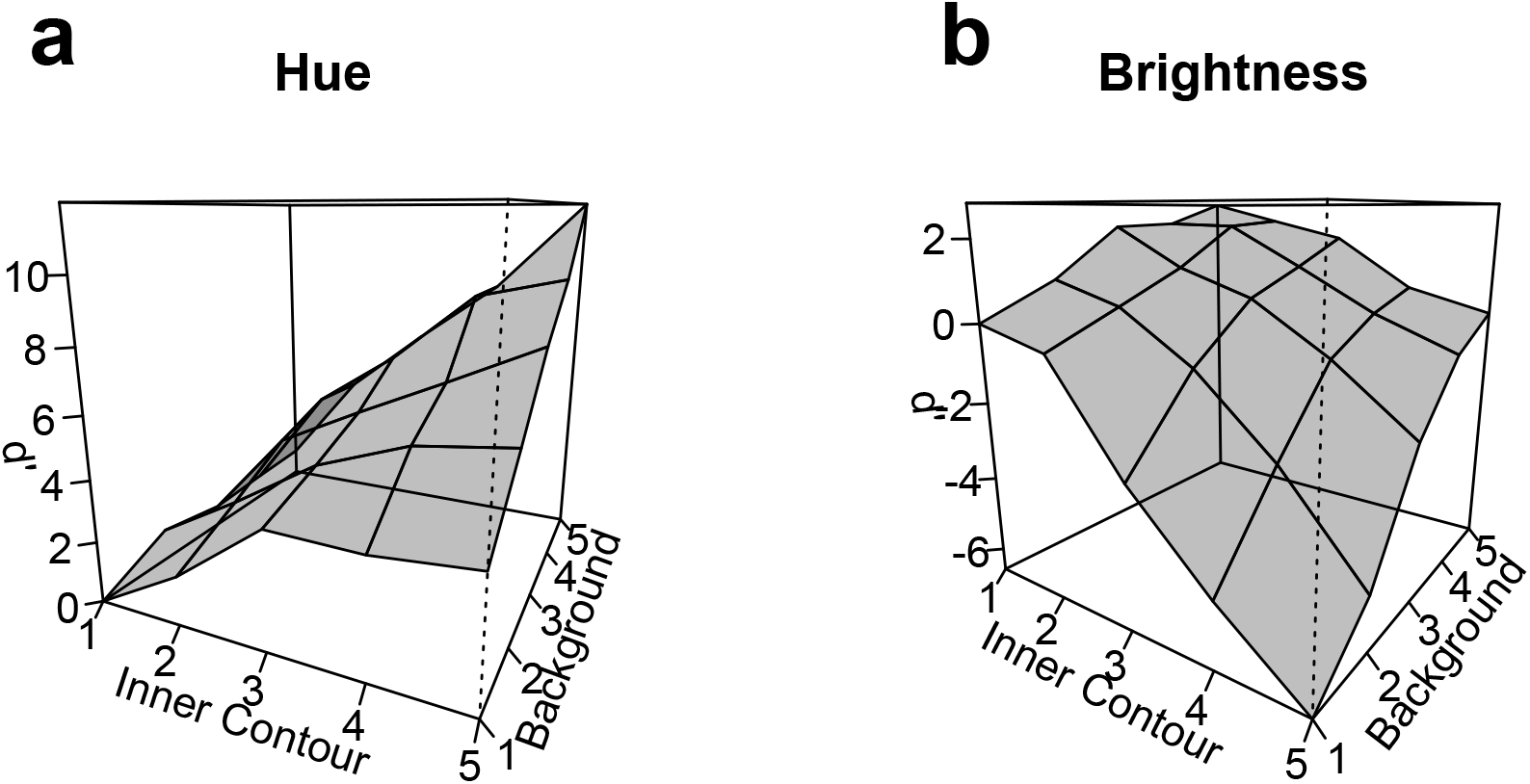
Three-dimensional surfaces for the average of the data over observers from Figure 7, showing the contributions of the inner contour and background to a) the hue and b) the brightness judgments under the saturated model.

**Table III:** Comparison of additive and saturated models for judgement based on hue

|  | Test stimuli (additive model × saturated model) |  |  |
| --- | --- | --- | --- |
|  | Df | Deviance | p-value |
| Obs. 1 | 16 | 105.44 | < 0.001 |
| Obs. 2 | 16 | 142.82 | < 0.001 |
| Obs. 3 | 16 | 104.81 | < 0.001 |
| Obs. 4 | 16 | 108.14 | < 0.001 |

**Table IV:** Comparison of additive and saturated models for judgement based on brightness

|  | Test stimuli (additive model × saturated model) |  |  |
| --- | --- | --- | --- |
|  | Df | Deviance | p-value |
| Obs. 1 | 16 | 56.154 | < 0.001 |
| Obs. 2 | 16 | 74.01 | < 0.001 |
| Obs. 3 | 16 | 32.83 | < 0.01 |
| Obs. 4 | 16 | 122.78 | < 0.001 |

## 4. Discussion

In the present study, MLCM was used to quantify the contributions to the filling-in strength of the WCE of two stimulus dimensions: the background and inner contour luminances. We quantified how changes in these features affected perceived filling-in using two separate response criteria, linked to the strength of the hue (the conventional WCE) and the brightness of the interior region. Control experiments using a stimulus contour that generated statistically undetectable filling-in confirmed that observers judged the perceived attributes of the interior regions and not changes in the background and contour luminances, per se. We also found that the results were largely independent of the ordering of presentation in a trial of both the backgrounds and the inner contours.

As found previously, the strength of the coloration effect depends on the luminance of the inner contour (Devinck & Knoblauch, 2012; Devinck et al., 2014; Gerardin et al., 2014). Here, we show that the stimulus configuration inducing the WCE generates both a hue and also a brightness filling-in of the interior area and that these two phenomena are differentially affected by the stimulus dimensions that we manipulated. The hue filling-in effect was assimilative, and the hue became more saturated with increases in luminance of the inner contour and the background. However, luminance of the interior contour generated a contrast effect for brightness, in that the judged brightness of the interior region decreased at higher contour luminances. As observers compared the interior regions of the two successively displayed stimuli and judged which central region appeared brighter, it is still possible that the perceived filling-in was assimilative, i.e., the same contrast polarity as the inner contour and that the effect of the contour luminance was simply to reduce the lightness. This is difficult to assess since we cannot simply compare the interior with the surround because both the surround and the interior vary in each condition. Additionally, if we hold the surround constant, then we cannot rule out its effect on the interior.

### The hue effect

Both the hue and the brightness of the interior region were judged to be greater with increases in the luminance elevation of the background. Pinna et al. (2001) have previously reported that color spreading in the WCE occurs not only with a white background but also with grey backgrounds and that even a faint spreading is perceived with dark backgrounds. Our results agree with these observations in that the background contribution to the hue judgments increased at higher luminances.

We also found that hue filling-in was more pronounced for bright than dim backgrounds. Previous studies have reported that color saturation diminishes when the brightness contrast between a colored object and its luminance background increases, a phenomenon named the gamut expansion effect (Brown & McLeod, ļ997) subsequently confirmed and quantified by several investigators (Bimler et al., 2009; Faul et al., 2008; Xing et al., 20ļ5). In contrast with these studies, our experiment shows that increasing the background luminance strengthens the assimilation hue, suggesting that we are observing a different phenomenon.

### The brightness effect

The WCE has typically been described as a coloration effect. Brightness variations in the interior region of the stimulus have not typically been systematically quantified. One exception to this concerns studies that used an achromatic stimulus configuration (Cao et al., 20ļļ; Coia & Crognale, 20ļ7). In Coia and Crognale (20ļ7), observers compared the filling-in region in the WCE to a reference stimulus with a physical luminance difference. Their matching results indicated that the test field shifted in the opposite direction from the inner contour showing an assimilation effect. In Cao et al. (2011), the luminances of the inner contour and the luminance of the background were fixed while the outer contour varied between high and low luminance levels, thus varying the contrast. Observers were asked to report which of two interior surface stimuli appeared darker. Their results followed a U-shape with the strength of the effect maximized for a range of medium luminance levels but not for the extreme luminance levels. Consequently, the luminance contrast between both contours affects the WCE but not linearly. An intriguing point is the authors’ description of their results: “Although there is an apparent assimilation effect in the chromatic WCE, it is hard to tell whether it is actually assimilation or some type of contrast effect happening here for the ‘opposite polarity’ condition”. As their method based the estimation of the WCE strength on the probability of discriminating which stimulus appeared darker, we can assume that if responses are inferior to 50%, then the surface appears lighter but not equal (which would be the case for responses equal to 50%), indicating a contrast rather than an assimilation effect. For our experiment based on brightness judgment, when the inner contour increases in luminance, the estimated contribution for this component decreases, indicating that the brightness of the interior region decreases, also indicating a contrast phenomenon. Thus, the two sets of results may be in agreement. Nevertheless, the nature of the brightness effect could be influenced by the presence of the chromatic component in our stimulus situation.

In Devinck et al. (2005), observers were allowed to adjust both the luminance and the chromaticity of a field in order to match the color of the WCE. The mean luminance match was near the luminance of the background, indicating that little or no luminance adjustment was required to make a perceptual match. The authors concluded that the WCE is predominantly a chromatic effect as originally suggested by Pinna et al. (200ļ). These results are not necessarily in conflict with the current study, in that we observed that the variation in brightness contrast with inner contour luminance is diminished at high backgrounds and might have been difficult to detect via matching. We would predict, then, that manipulating the luminance of the background would affect the luminance match, but this would require a more systematic study of the background than was performed by Devinck et al. (2005). Here, our data indicate that the perceptual effect is not limited only to a coloration phenomenon in that both the background and inner contour luminances influence observers’ judgment of the brightness of the central surface. A simple hypothesis to account for the reduced brightness effect at high background luminances is to suppose that the background light added to the interior region cancels the contrast effect. This would not explain, however, the fact that higher background luminances led to a stronger hue percept.

### Assimilation vs contrast

Whether one observes a contrast or assimilation effect may depend on the width of the contours (Fach & Sharpe 1986; Helson & Rohles, 1959; Helson, 1963). Other factors such as the luminance of the inducing stimuli can also influence our percept. Thus, de Weert and Spillmann (1995) indicated that assimilation or contrast occurred depending on whether the inducing contours of varied reflectance were darker or lighter than the gray background in using a pincushion pattern. A matching experiment for brightness judgements indicated that contrast occurred when the luminance level of the inducing contour was above the luminance level of the background and that assimilation occurred when the luminance of the inducing contour was below the luminance of the background. Given the different spatial dependencies of chromatic and luminance sensitive mechanisms, one might expect differences in the spatial domains over which chromatic and luminance components of a stimulus induce assimilation and contrast. Given the dependence of induction phenomena on stimulus configuration, however, (Fach & Sharpe 1986; de Weert & Spillmann 1995; Smith et al. 2001; Monnier & Shevell 2003, 2004), it is difficult to predict *a priori* whether the dimensions of the contours that we used should predict one or the other for the hue and brightness judgments.

In our study, observers compared the interior regions of the two successively displayed stimuli and judged which central region appeared brighter. Under this condition, our results indicated that the brightness of the interior region decreases generating a relative contrast rather an assimilation effect. However, it still possible that different visual phenomena are perceived in other circumstances. In making a brightness judgements of the central region with respect to the outer region, we noted an assimilation phenomenon at the lowest luminance level of the background and at the highest luminance level of the inner contour. This condition corresponds to the lower left corner in Figure 2.

### Unitary vs multiple mechanisms

The color of the central surface in the WCE is characterized by a spread of color from the inner contour. Most previous studies of the WCE reported that the coloration effect depends on both the chromatic and luminance contrasts of the inner and outer contours. For example, most authors demonstrated that the coloration effect increases with increasing luminance contrast between inner and outer contours (Devinck & Knoblauch, 20ļ2; Devinck et al., 2005). Additionally, the coloration effect increases when the chromatic coordinates of the inner and outer contours are approximately complementary in the color diagram (Devinck et al., 2006). Thus, the main explanation assumes a filling-in process in which a neuronal mechanism detects the contour and generalizes it beyond the confines of the immediate stimulus. Most studies in the WCE have reported an important role for several types of contour mechanisms generating a long-range filling-in percept. Taken together, these data suggest that the filling-in process involved in the WCE requires multiple levels of processing (Devinck et al., 2014b; Pinna et al., 2001; Pinna & Grossberg, 2005; von der Heydt & Pierson, 2006). The present results indicate that the mechanisms inducing the brightness and coloration percept in the WCE are affected differently by the luminance of the inner contour. These opposing responses due to the inner contour suggest that multiple mechanisms contribute to the appearance of the interior region. Different mechanisms could be activated or inhibited yielding to color assimilation or brightness contrast effects, respectively. Future experiments based on visual masking or contour adaptation could investigate such phenomena.

## Acknowledgements

This work was supported by a grant from the Agence Nationale de la Recherche to FD (ANR-11-JSH-20021) and by LABEX CORTEX (ANR-11-LABX-0042).

